# Prospective real-time metagenomic sequencing during norovirus outbreak reveals discrete transmission clusters

**DOI:** 10.1101/473405

**Authors:** Amanda M. Casto, Amanda L. Adler, Negar Makhsous, Kristen Crawford, Xuan Qin, Jane M. Kuypers, Meei-Li Huang, Danielle M. Zerr, Alexander L. Greninger

**Author notes:** Corresponding Author: 1616 Eastlake Avenue East, Suite 320 Seattle, WA 98102. Alternative corresponding author: Amanda Casto 1616 Eastlake Avenue East, Suite 320 Seattle, WA, 98102. These authors contributed equally to this manuscript (Amanda M. Casto, Amanda L. Adler).

## Abstract

**Background:** Norovirus outbreaks in hospital settings are a common challenge for infection prevention teams. Given the high burden of norovirus in most communities, it can be difficult to distinguish between on-going in-hospital transmission of virus and new introductions from the community and challenging to understand the long-term impacts of outbreak-associated viruses within medical systems using traditional epidemiological approaches alone.

**Methods:** Real-time metagenomic sequencing during an on-going norovirus outbreak associated with a retrospective cohort study.

**Results:** We describe a hospital-associated norovirus outbreak that affected 13 patients over a 27-day period in a large tertiary pediatric hospital and was chronologically associated with a spike in self-reported gastrointestinal symptoms among staff. Real-time metagenomic next-generation sequencing (mNGS) of norovirus genomes demonstrated that 10 chronologically overlapping hospital-acquired norovirus cases were partitioned into three discrete transmission clusters. Sequencing data also revealed close genetic relationships between some hospital-acquired and some community-acquired cases. Finally, this data was used to demonstrate chronic viral shedding by an immunocompromised hospital-acquired case patient. Analysis of serial samples from this patient provided novel insights into the evolution of norovirus within an immunocompromised host.

**Conclusions:** This study documents one of the first applications of real-time mNGS during a hospital-associated viral outbreak. Given its demonstrated ability to detect transmission patterns within outbreaks and elucidate the long-term impacts of outbreak-associated viral strains on patients and medical systems, mNGS constitutes a powerful resource to help infection control teams understand, prevent, and respond to viral outbreaks.

**Summary Statement:** Real-time metagenomic sequencing performed during a hospital-associated norovirus outbreak identified genetically-distinct, chronologically-overlapping case clusters. After the epidemiologically-defined outbreak had ceased, on-going transmission and shedding of outbreak-associated virus was also detected. These findings illustrate the value of genomics as a tool for infection control.

## Introduction

Norovirus is the most common cause of acute gastroenteritis in the United States with an estimated incidence of 19 - 21 million cases per year [1]. The high prevalence of norovirus complicates efforts to prevent introduction into health care settings from the community [2]. Transmission primarily occurs through the fecal-oral route, though it can also occur via aerosolized viral particles and environmental contamination [3,4]. Most cases of norovirus resolve in 12 - 60 hours, but hospital-associated outbreaks frequently involve immunocompromised patients, who can experience prolonged symptoms and viral shedding [5–8] and potentially act as reservoirs for viral transmission.

Metagenomic next-generation sequencing (mNGS) is increasingly used for detection of infectious organisms when other clinical testing has been non-diagnostic. The same technique can be employed to elucidate pathogen transmission chains [9–11]. We and others have previously used real-time metagenomics to both rule-in and rule-out on-going transmission of hospital-acquired respiratory virus infections [12–14].

Here we describe a hospital-associated outbreak of norovirus that affected 13 patients over 27 days in a large pediatric hospital and was temporally associated with a spike in self-reported gastrointestinal symptoms among staff. We discuss the use of real-time mNGS to identify transmission clusters within the outbreak and to elucidate the long-term impacts of the outbreak including on-going transmission of an outbreak strain after hospital-acquired cases had ceased and chronic shedding of hospital-acquired virus in an immunocompromised host.

## Methods

### Setting

Seattle Children’s Hospital (SCH) is a 354-bed tertiary-care pediatric facility located in Seattle, Washington. The medical unit is a 64-bed unit located on two floors that cares for patients with a wide variety of acute and chronic health issues. The cancer care unit is a 48-bed unit located on two floors that cares for patients with hematologic and oncologic diagnoses, including those undergoing hematopoietic cell transplant (HCT). These two units are in the same building of the hospital. All rooms on these units are private rooms with en-suite restrooms.

### Patient and staff case definitions

Hospital-acquired norovirus cases were hospitalized patients who developed acute gastrointestinal symptoms (vomiting, abdominal pain, diarrhea, or nausea) more than 48 hours after admission or within 24 hours of discharge and had a positive norovirus RT-PCR test [15] (see Supplementary Methods). Staff cases included any hospital staff member with self-reported sudden-onset gastrointestinal symptoms.

### Case detection

In January 2017, a routine daily safety briefing revealed that seven staff members assigned to the medical unit had called in sick due to gastroenteritis symptoms within a 3-day period. During initial discussions with unit staff, four patients were also identified with hospital-acquired gastroenteritis with similar dates of symptom onset. Stool RT-PCR testing confirmed that all four patients had norovirus. The subsequent investigation found delays in initiating isolation precautions and revealed interactions between some ill patients and staff members. It was ultimately discovered that a recent change in the name of the norovirus test resulted in the exclusion of norovirus results from the electronic report used by the Infection Prevention Department to perform routine hospital surveillance. Therefore, a list of patients with norovirus-positive testing was obtained from the clinical laboratory and a retrospective review was performed. This review identified two additional hospital-acquired norovirus cases on the cancer care unit that occurred 4 and 7 days prior to the first case on the medical unit, respectively. The first of these cases was considered the start of the outbreak (Day 0). A communication was sent to providers and nurses requesting that patients with hospital-acquired gastroenteritis be tested for norovirus. The occupational health department monitored staff members with gastroenteritis symptoms and enforced work restrictions. By Day 33 (three incubation periods since the last case), no further hospital-acquired cases were identified and the outbreak was considered resolved. Measures performed to control the outbreak are described in the Supplementary Methods.

### mNGS library generation and sequencing

Prospective mNGS aimed at whole genome recovery was performed on Days 9 and 26 of the outbreak. We sequenced samples from the hospital-acquired norovirus cases and from staff cases who tested positive for norovirus. To understand the diversity of strains circulating in the community, samples collected from patients at facilities within the University of Washington (UW) medical system, including SCH patients with community-acquired norovirus, were also sequenced (Supplementary Table S1). mNGS sequencing was performed as described previously [12,16] (see Supplementary Methods).

### Genotype Assignment

Sequencing reads were adapter and quality trimmed using cutadapt and de novo assembled using SPAdes v3.11 [17,18]. The resulting scaffolds were used as input for the online norovirus genotyping tool [19]. There were representatives of 4 genotypes among our sequences. A reference sequence was selected from GenBank for each of these genotypes: ORF1: GII.P16, ORF2: GII.4, Sydney_2012 (subsequently referred to as the “Sydney” genotype) – LC175468 [20]); ORF1: GII.P16, ORF2: GII.2 (“GII.2” genotype) – KY905336; ORF1: GII.P12, ORF2: GII.3 (“GII.3” genotype) – KY905334; ORF1: GII.P7, ORF2: GII.6 (“GII.6” genotype) – KU935739.

### Phylogenetic and Evolutionary Analysis

Consensus sequences were generated for each sequenced sample by mapping reads to the corresponding reference sequence using Geneious v10 [21]. For each sequence, single nucleotide variants (SNVs) relative to the reference were called by manual review (see Supplementary Methods). Sequences were aligned using MAFFT v7.222 with the default settings [22]. Phylogenic trees were generated using MrBayes with chain length 1,100,000 [23]. dN/dS analyses were performed on an alignment of sequences from Case 2.

### Sequence Clusters

Two sequences were said to belong to a cluster if they had 10 or fewer SNV differences. A sequence would be added to an existing cluster if it differed from at least one sequence in the cluster by 10 SNVs or less. The selection of 10 differences to define a cluster was based on past estimations of the mean norovirus genome evolutionary rate, corresponding to a time to most recent common ancestor of approximately 6 weeks, and on the distribution of pairwise differences between the sequences (see Supplementary Methods).

### Results

### Outbreak characterized by chronologically overlapping patient and staff cases

Over a period of 27 days, there were 13 confirmed hospital-acquired norovirus cases (Figure 1). Eight of these were on the medical unit and five were on the cancer care unit (Supplementary Notes S1 and S2). Over the same time period, 86 staff members self-reported gastrointestinal symptoms to occupational health, though only 16 (19%) of these individuals were known to be assigned to one of the affected units (medical = 9 and cancer care = 7). Norovirus testing was performed on only two staff members and both were confirmed to be norovirus positive.

**Figure 1:**
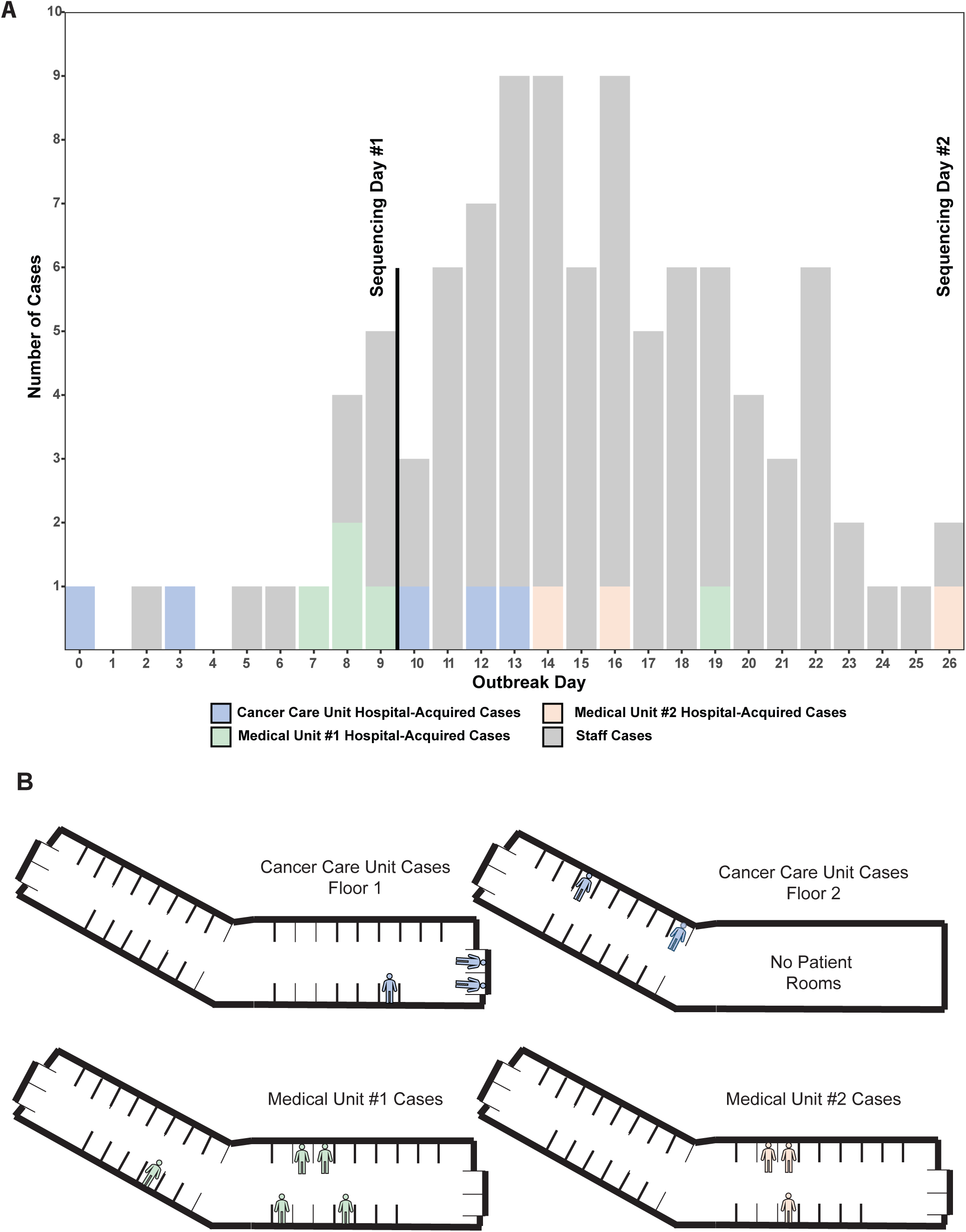
A) Epidemiologic curve including hospital-acquired norovirus cases by hospital unit and staff with gastrointestinal symptoms. B) Diagram showing location of hospital-acquired norovirus cases within the hospital.

### Hospital-acquired cases phylogenetically cluster according to hospital geography

We were able to sequence samples for 10 of the 13 patient cases, six on the first day of prospective sequencing (Day 9) and four on the second (Day 26). Phylogenetic analysis of these sequences revealed 3 distinct clusters corresponding to hospital geography with one on the cancer care unit (Cancer Care Unit Cluster) and one each on two different floors of the medical unit (Medical Unit Clusters #1 and #2) (Figures 2 and 3). The 6 samples sequenced early in the outbreak were divided between the Cancer Care Unit Cluster and Medical Unit Cluster #1 (Supplementary Figure S1) and were all members of the Sydney genotype. The 4 samples sequenced later in the outbreak belonged to the Cancer Care Unit Cluster and Medical Unit Cluster #2 (Supplementary Figure S1). Samples in the Medical Unit Cluster #2 were of the GII.2 genotype. Interestingly, four samples from the Cancer Care Unit Cluster were genetically identical even though they were collected up to 18 days apart (Supplementary Table S2).

**Figure 2:**
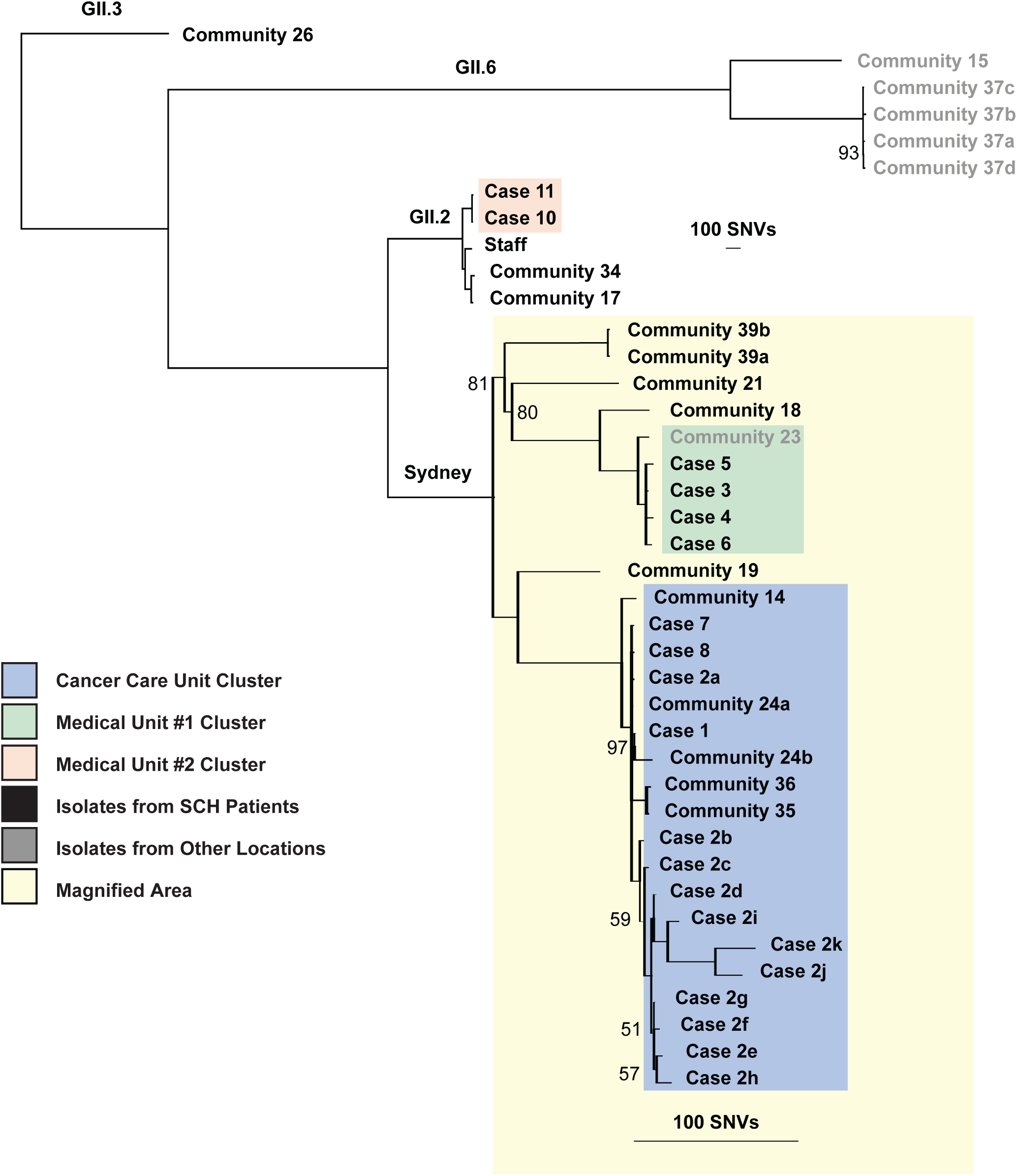
Phylogenetic tree of hospital- and community-acquired norovirus sequences. Colored boxes represent hospital-acquired case clusters. Black lettering represents samples collected at SCH while gray lettering represents samples collected at other facilities. All non-100% posterior probabilities are denoted.

**Figure 3:**
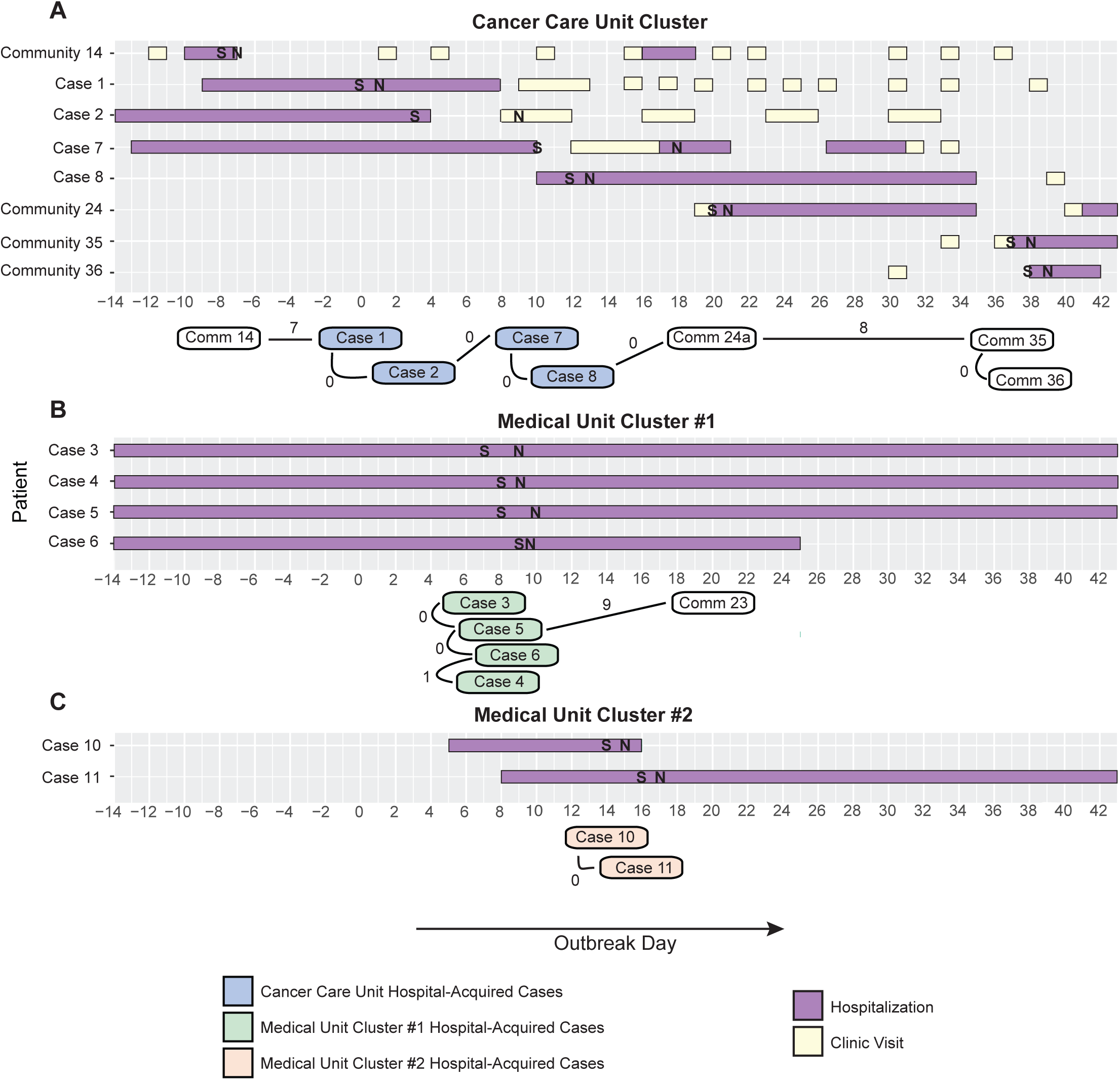
The three transmission clusters within the norovirus outbreak. Timelines of gastrointestinal symptoms, norovirus testing, hospitalizations, and outpatient visits for pediatric cluster members are shown on the top of each panel. The number of SNV differences between cluster members is shown on the bottom of each panel.

On the second day of sequencing (Day 26), we were also able to generate a sequence for one of the two staff members with confirmed norovirus. This sample represented the GII.2 genotype but was quite distinct from the two GII.2 patient cases (117 and 118 SNV differences, respectively).

For each of the two rounds of sequencing our turn-around time from samples to phylogenetic analysis was approximately 24 hours with information about the genetic relationships among the hospital-acquired patient and staff cases relayed to the infection prevention team in real-time.

### Multiple putative cryptic transmissions noted between community-acquired and hospital-acquired cases

We also generated sequences for a total of 19 community-acquired norovirus infections in SCH patients and patients at other UW medical facilities that were collected during and in the months after the outbreak. These samples represented 4 different norovirus genotypes. Most were genetically distinct from one another and from the 3 hospital-associated transmission clusters. There were, however, a total of 4 community-acquired infections in SCH patients that grouped with the Cancer Care Unit Cluster (Supplementary Figure S1). One of these samples was from a patient diagnosed just prior to the outbreak; the remaining 3 infections were diagnosed after the outbreak. All 4 infections were in oncology patients who were found to have shared clinic or housing space with hospital-associated case patients (Figure 3). Interestingly, a sample from an adult patient who was hospitalized at another facility during the outbreak fell within Medical Unit Cluster #1.

### Hospital-acquired cases were often persistently positive on follow-up testing

Subsequent norovirus testing was performed on 6 of the 13 (46%) case patients in the 9 months following the outbreak and 5 (83%) remained positive. These included 2 patients from the medical unit (Cases 3 and 4) and 3 patients from the cancer care unit (Cases 2, 7, and 9). Positive follow-up samples were collected a median of 57 days after the respective outbreak sample (range, 9-258 days).

### Follow-up testing reveals long-term shedding in an immunocompromised patient

The second hospital-acquired norovirus case (Case 2) was a 2-year old who had received an HCT for hemophagocytic lymphohistiocyotosis (HLH) that was complicated by graft versus host disease. The patient had a total of 8 samples collected during and in the first 5 months after the outbreak (Case 2a-h). The patient’s blood cell counts were stable and immunosuppressive medications were not significantly changed during this time. The patient then experienced reactivation of HLH and was treated with 3 months of chemotherapy, during which sample Case 2i was collected. Finally, the patient underwent a second HCT, after which samples Case 2j and Case 2k were collected. We were able to generate sequences for all 11 samples collected from this patient.

All Case 2 samples were of the same genotype and the genetic distance between samples gradually increased with time between sample collection, suggesting chronic infection rather than resolution and re-infection. Over a period of 258 days, this patient’s norovirus accrued 65 consensus sequence changes relative to the first sample with approximately equal numbers of synonymous (33) and non-synonymous (32) changes (Supplementary Figures S3 and S4, Supplementary Table S3). The number of non-synonymous changes was greater than the number of synonymous changes in all samples except for the last one, Case 2k. The rate of accumulation of consensus changes seems to have increased slightly after the patient experienced recurrence of their HLH, received chemotherapy, and underwent a second HCT.

A number of non-synonymous changes were noted in the P2 subdomain of ORF2, which encodes the most immunogenic part of the norovirus capsid and is thought to be important to immunologic escape by viruses [24]. The average value of dN/dS (when Case 2b-k were compared to Case 2a) for ORF2 was 1.717 (standard deviation 0.394). The patient’s ORF2 dN/dS was substantially larger than a previous dN/dS estimate of 0.12 for GII.4 ORF2 sequences [25]. We were unable to calculate dN/dS for the P2 subdomain as there no synonymous consensus changes observed in this region for any of the later samples relative to Case 2a. The number of non-synonymous consensus changes observed ranged from 1 to 8.

The average mutation rate in ORF2 for the same comparisons was 0.015 substitutions/site/year (standard deviation 0.011), which again was substantially higher than GII.4 ORF2 mutation rate estimates of 5.4 × 10^−3^ substitutions/site/year [25].

## Discussion

We describe a complex hospital-associated norovirus outbreak analyzed with prospective and retrospective mNGS with a sample to phylogenetic analysis turn-around time of 24 hours for prospective sequencing.

Through the application of genomic analyses to this outbreak, we gained new insights into viral transmission among case patients and into the relationship between hospital-acquired and community-acquired cases. Firstly, we showed that what appeared to be one large, hospital-wide norovirus outbreak was composed of three genetically distinct transmission clusters, implying the introduction of three different norovirus strains into the hospital within a short period of time. This finding highlights the importance of health care workers staying home when ill and of screening hospital visitors for symptoms of viral illness. Transmission of these strains within the hospital demonstrates the importance of hand hygiene, environmental cleaning, and prompt isolation of symptomatic patients, which can be challenging in patient populations that have many potential noninfectious etiologies for vomiting and diarrhea.

Secondly, we found that there were no SNV differences among cases on the cancer care unit, suggesting to us that a contaminated fomite may have propagated the outbreak on this unit rather than person-to-person transmission. While environmental surfaces were cleaned with bleach during the outbreak, this finding suggests that efforts should have been intensified on this unit.

Our analysis also showed a close relationship between the cancer care unit cases and several community-acquired cases in other pediatric oncology patients. This finding suggests that the cancer care unit viral strain may have continued to be transmitted after the epidemiologically-defined end of the outbreak. A possible venue for these putative transmission events was the outpatient oncology clinic, highlighting the potential for viral spread in ambulatory settings, a particular concern for immunocompromised patients.

Finally, we showed that the noroviruses responsible for the outbreak were reflective genetically of the noroviruses circulating in the community at the time. In particular, both of the genotypes observed among hospital-acquired cases were seen in community-acquired cases. More broadly, our genetic analysis illustrates how outbreaks are dependent on pathogen circulation in the community. The existence of three discrete transmission clusters in this outbreak required that there was genetic diversity among viruses in the community and was also likely reflective of a high overall burden of norovirus locally at the time of the outbreak, which took place in January, the typical peak of norovirus season [26].

The above findings demonstrate the utility of genetic analyses in outbreak investigations. Indeed, some of these findings would not have been apparent without the single nucleotide resolution provided by whole genome sequencing. We have shown that mNGS can be performed in a timely manner comparable to genotyping assays. In addition to the high resolution genomic data they produce, mNGS approaches have the advantage of being easy to adapt for other organisms, including those for which little genomic information is available. Furthermore, data from mNGS analyses can be directly compared to data from other studies and can be pooled across studies for new analyses.

Our genetic analysis of this outbreak also gave us the opportunity to examine norovirus evolution in an immunocompromised patient who was one of the hospital-acquired cases. We found that this patient’s virus accumulated non-synonymous mutations in ORF2 and particularly in the P2 subdomain at a faster rate than synonymous mutations. This same pattern was found when examining genomes from other chronically infected, immunocompromised patients, but was not seen when the evolution of GII.4 genotype viruses at large was examined (Supplementary Note S3, Supplementary Table S4) [25]. We hypothesize that the norovirus population size in immunocompromised patients tends to be larger due to their weakened immunity. Population size is also likely more stable than in immunocompetent individuals as norovirus that “resides” in an immunocompromised host does not experience the bottlenecks associated with transmission from host to host. The net effects of these differences in population dynamics is that genetic drift, which can drive the fixation of neutral synonymous mutations, is weaker and that positive selection is more effective (as advantageous non-synonymous mutations are unlikely to be lost due to drift) in immunocompromised hosts, even in the context of weakened selective pressures from the immune system. This is one possible explanation for both the surplus of non-synonymous mutations and the deficit of synonymous mutations that we observed in the ORF2 gene and P2 subdomain in samples from the Case 2 patient.

In some other viral species, viruses that evolve in immunocompromised hosts seem to predict broader evolutionary trends [27]. It is currently unknown if noroviruses from immunocompromised hosts influence viral evolution or even if viral transmission from chronically infected patients is possible. It seems unlikely, though, that viruses from immunocompromised hosts have no impact on norovirus evolution given that there are many degrees of immunocompromise, that the number of immunocompromised individuals continues to increase, and that immunocompromised patients have frequent contact with the health care system and each other.

This study was limited by our inability to obtain specimens from ill staff, a challenging problem as most otherwise healthy patients do not seek medical attention for norovirus infection and hospital staff often seek care from outside providers. A further limitation was the study’s mostly retrospective nature. While the use of Illumina sequencing allows for high-quality deep sequencing, the cost of reagent cartridges forces labs to use them in batched sequencing processes, which somewhat limited our ability to perform a true “real-time” analysis [28,29]. Finally, our failure to recover genomes from some samples illustrates the challenges of working with clinical samples with variable viral RNA quality and quantity using mNGS compared with other methods such as capture sequencing that require longer turn-around times [30,31]. Stool is a particularly daunting sample source given the number and diversity of other organisms present [32].

Overall, our results demonstrate how metagenomics methods can be employed in outbreak investigations to elucidate intricate details about pathogen transmission. This information is vital to efforts aimed at optimizing infection control interventions [26]. Our findings underscore the critical importance of these interventions in both inpatient and ambulatory settings to prevent the transmission of pathogens in health care facilities. Infectious control efforts are perhaps particularly crucial for norovirus, a common and highly contagious organism that can have lasting impacts on medical systems and on vulnerable patient populations [33].

## Acknowledgements

We thank the patients and staff of Seattle Children’s Hospital and the University of Washington Clinical Virology Laboratory for their contributions to the outbreak response and clinical testing described here.

## Funding

This work was supported by departmental start-up funds from the Department of Laboratory Medicine at the University of Washington.

## Conflict of Interest Statement

The authors have no conflicts of interest to declare.

